# Local nutrient hotspots shape red deer habitat selection in a nutrient-poor landscape

**DOI:** 10.64898/2026.08.17.745224

**Authors:** Elke Wenting, Manon van den Braak, Chris Vervoorn, Henk Luten, Ruben Vermeer, Dennis R. Lammertsma, Lysanne Snijders, Elisabeth S. Bakker, Andrea Kölzsch

**Affiliations:** Biodiversity Research Institute (CSIC – University of Oviedo – Principality of Asturias), 33600 Mieres, Spain; Radboud University, Radboud Institute for Biological and Environmental Sciences, Department of Ecology, Box 9010, 6500 GL Nijmegen, the Netherlands; Netherlands Institute of Ecology (NIOO-KNAW), Department of Terrestrial Ecology, Droevendaalsesteeg 10, 6708 PB Wageningen, the Netherlands; Behavioural Ecology Group, Wageningen University & Research, De Elst 1, Wageningen, 6708 WD, The Netherlands; Wildlife Dierenarts Luten, Alteveerselaan 30a, 6881 AW Velp, The Netherlands; Natuurmonumenten, Beheereenheid Veluwezoom, Heuvenseweg 6a, 6991 JE Rheden, the Netherlands; Wageningen University & Research, Wageningen Environmental Research, Droevendaalsesteeg 3a, 6708 PB Wageningen, The Netherlands; Wageningen University, Wildlife Ecology & Conservation Group, Droevendaalsesteeg 3a, 6708 PB Wageningen, the Netherlands; Max Planck Institute of Animal Behavior, Department of Migration, Am Obstberg 1, 78333 Radolfzell, Germany

**Keywords:** habitat selection, large herbivores, nutrient heterogeneity, nutrient redistribution, selective feeding

## Abstract

Nutrient availability in many temperate ecosystems is shaped by soil properties and historical land use. Especially in otherwise nutrient-poor landscapes human-induced, local fertilisation can generate fine-scale mosaics of nutrient hotspots. Whether and how large herbivores respond to such heterogeneity remains poorly understood. We tested whether spatial variation in soil-derived nutrient availability structures habitat selection by large herbivores, using full-year GPS tracking data from 7 red deer (*Cervus elaphus*) in the Veluwe, the Netherlands. We used soil types as a proxy for nutrient availability and assigned nutrient scores based on soil pH, cation exchange capacity and soil structure. We then evaluated habitat selection across multiple components of space use: (i) home range size; (ii) use of relatively nutrient-rich parts within home ranges; (iii) selection among soil types; and (iv) selection of locally enriched former agricultural patches. Red deer used relatively nutrient-rich within their home range more than expected based on availability, including local patches enriched by former agricultural use. However, site selection did not consistently follow nutrient scores among soil types. These results show that nutrient availability does shape habitat selection, but primarily through fine-scale, localised nutrient enrichment rather than broad-scale variation in soil properties. Our findings demonstrate that nutrient-related foraging contributes to habitat selection in a large wild herbivore, while also revealing that this process is scale- and context-dependent. By repeatedly concentrating their foraging in nutrient-rich patches, large herbivores may contribute to nutrient redistribution across the landscape, with the potential to reinforce or modify existing spatial heterogeneity in resource availability and ecosystem functioning.

## 1. Introduction

Nutrient availability is spatially heterogeneous in many temperate landscapes, shaped by variation in soil properties, historical land use and human-induced inputs such as long-term atmospheric nitrogen (N) deposition (e.g. Song et al. 2023; Vogels et al. 2023; Van der Plas et al. 2024). Such heterogeneity can create mosaics of resource quality that influence how animals move, forage and use space, in order to acquire the full spectrum of chemical elements that are required for growth, reproduction, and survival (Kaspari 2021).

Large herbivores play a central role in shaping vegetation structure, competitive interactions among plant species and nutrient cycling (e.g. Riesch et al. 2022; Pringle et al. 2023; Garcia et al. 2025). The spatial pattern of these effects is driven by their foraging movement, which may depend on whether herbivores respond to underlying gradients in nutrient availability by concentrating their foraging activity in relatively nutrient-rich areas. Such spatially structured herbivory may, in turn, reinforce heterogeneity in vegetation dynamics and other herbivore-mediated ecosystem functions such as nutrient cycling (e.g. Garcia et al. 2023; 2025).

However, most studies of habitat use by herbivores derive resource quality from vegetation structure or land-cover categories (e.g. Fricke et al. 2022; Rech et al. 2025), rather than directly accounting for underlying variation in soil properties. This limits our ability to evaluate resource availability in herbivore-driven ecological processes, such as selective feeding for nutrients (Raubenheimer et al. 2009; Raubenheimer & Simpson 2016). Studies from African savannas have shown that herbivores preferentially use nutrient-rich hotspots created by former livestock corrals (Augustine 2003; Augustine et al. 2003), demonstrating that spatial variation in soil fertility can influence habitat use across diverse ecosystems. However, such studies have largely focused on discrete nutrient hotspots, whereas less is known about how large herbivores respond to broader gradients in nutrient availability across heterogeneous landscapes. A better understanding of such feeding strategies requires explicitly linking spatial patterns of herbivore movement to heterogeneity in nutrient availability that are independent of vegetation structure.

Particularly in nutrient-poor landscapes, nutrient availability is often highly uneven, with some patches more depleted or enriched than others. Relatively nutrient-rich patches are typically distributed heterogeneously due to both intrinsic soil variation and historical land use (Abadie et al. 2018). Herbivores may adapt their movement patterns and foraging sites to such spatial variation (Martin et al. 2015). Such responses are likely to occur across multiple spatial scales, from differences in home-range size to selection among soil types and fine-scale use of locally enriched patches such as former agricultural fields (Hjermann et al. 2025). Multi-scale responses would indicate that nutrient heterogeneity is an important driver of structuring the spatial distribution of herbivore pressure, which may be even more evident under nutrient-poor conditions. Here, scarcity of nutrients may both result from absolute low levels of nutrient availability or from nutrient imbalances, affecting the relative abundance of nutrients needed for herbivore physiology.

Empirical evidence for the hypothesis that animals adapt their movement patterns to soil heterogeneity remains limited, particularly at landscape scales and for free-ranging animals. While studies of diet selection and controlled experiments demonstrate that herbivores respond to nutrient availability (e.g. Ginane et al. 2005; Blubaugh 2023), it remains unclear how these responses translate into spatial patterns of habitat use in natural systems.

Here, we investigate how soil types, as a proxy for spatial variation in nutrient availability, influence patterns of habitat use by a large herbivore in a nutrient-poor ecosystem. We use GPS tracking data of red deer (*Cervus elaphus*) in the Veluwe, the Netherlands, a landscape characterised by nutrient-poor sandy soils, long-term N deposition, and heterogeneity in soil properties and locally abandoned former agricultural fields (e.g. Siepel et al. 2018; Vogels et al. 2024). We hypothesised that spatial variation in soil-derived nutrient availability structures habitat selection by large herbivores under generally nutrient-poor conditions, which enables them to fulfil their nutritional demands. We predicted that (i) individuals in more nutrient-poor areas maintain larger home ranges; (ii) individuals use the most nutrient-rich areas within their home ranges; (iii) individuals select to forage more on the relatively nutrient-rich soil types within their home ranges; and (iv) individuals select locally enriched patches associated with former agricultural land use more than surrounding areas, reflecting fine-scale and local responses to nutrient heterogeneity.

## 2. Methods

### 2.1 Study area

This study was conducted in the southeastern part of the Veluwe (52°02’N, 6°01’E), the Netherlands, covering an approximately 100-200 km² landscape of forests, heathlands, grasslands, and former agricultural fields. The area mainly consists of glacier deposits and cover sands, but was once an agrosilvopastoral landscape (Kuiters 2005). Habitat types scattered around the landscape include dry grass-heathlands, pastures, abandoned crop fields and different kinds of woodland (Kuiters 2005).

The sandy soils of the Veluwe are generally nutrient-poor and acidic, conditions that have been further influenced by long-term atmospheric N deposition and associated acidification (e.g. Song et al. 2023; Vogels et al. 2023; Van der Plas et al. 2024). These conditions, that have been demonstrated to reduce the fitness, reproduction and survival in different trophic levels, cause imbalances in elemental stoichiometry (e.g. Briggs & Mainwaring 2022; Vogels et al. 2024), which makes the area suitable for examining whether red deer respond to spatial variation in soil-related nutrient availability under nutrient poor conditions.

Part of the area inhabits free-ranging Scottish Highland cattle (*Bos taurus*) and Icelandic horses (*Equus caballus*), introduced in the 1980s, aiming to stimulate natural processes with minimised human intervention (Groot Bruinderink & Lammertsma 2001; Kuiters 2005). Other free-ranging ungulates in the area include red deer, fallow deer (*Dama dama*), roe deer (*Capreolus capreolus*) and wild boar (*Sus scrofa*). Large carnivores such as wolves (*Canis lupus*) were absent at the time of GPS-tracking. Also, as part of the management policy, there is no supplementary feeding of herbivores, meaning that active control of area use is not applicable (Groot Bruinderink et al. 2000).

### 2.2 Study species

Red deer are intermediate feeders, combining grazing and browsing depending on seasonal resource availability and habitat conditions (Krojerová-Prokešová et al. 2010). Their daily food intake averages 4.5-5 kg/day of dry matter for males and 3-3.5 kg/day for females, although it varies with age, sex and season (e.g. Kay & Staines 1981; Putman 2012). Home range sizes are influenced by many environmental and individual factors, including season, calving, hunting pressure, landscape type, mating season, and sex (Reinecke et al. 2014; Jarnemo et al. 2023), and are generally smaller during winter-spring, calving, and hunting periods (Georgii & Schröder 1983; Rivrud et al. 2010; Jarnemo et al. 2023), as well as in forested areas compared to mixed habitats (Borkowski et al. 2016), and females generally had smaller ranges than males (Jarnemo et al. 2023).

### 2.3 GPS data collection and home range calculation

Four female and three male red deer of varying ages were fitted with GPS collars (Vectronics GPS Plus collar) during the years 2011 and 2015 (Fig. 1), which recorded their location once per hour, with tracking durations varying between individuals (Table 1). This animal experiment was conducted under Dutch law (The experiments on Animals Act; Wet op de dierproeven) after approval by the ethics committee (DEC ALT 09.03). Also a license was obtained concerning the Flora and Fauna Act; Flora en faunawet; license FF/75A/2009/031). Red deer were captured using a dart gun using anaesthesia (Large Animal Immobilion ®Novartis). After collaring, the anaesthesia was reversed using an antidote (Large Animal Revivon ®Novartis). To ensure that the anaesthesia used on the deer does not affect their spatial behaviour, the two weeks after fitting the GPS collar are excluded from the analyses. At the moment of capturing, blood samples were taken for the assessment of possible contagious diseases and genetic research, that we also analysed for elemental concentrations as contextual information (see Supplementary Information, Fig. S1-S3).

**Figure 1.**
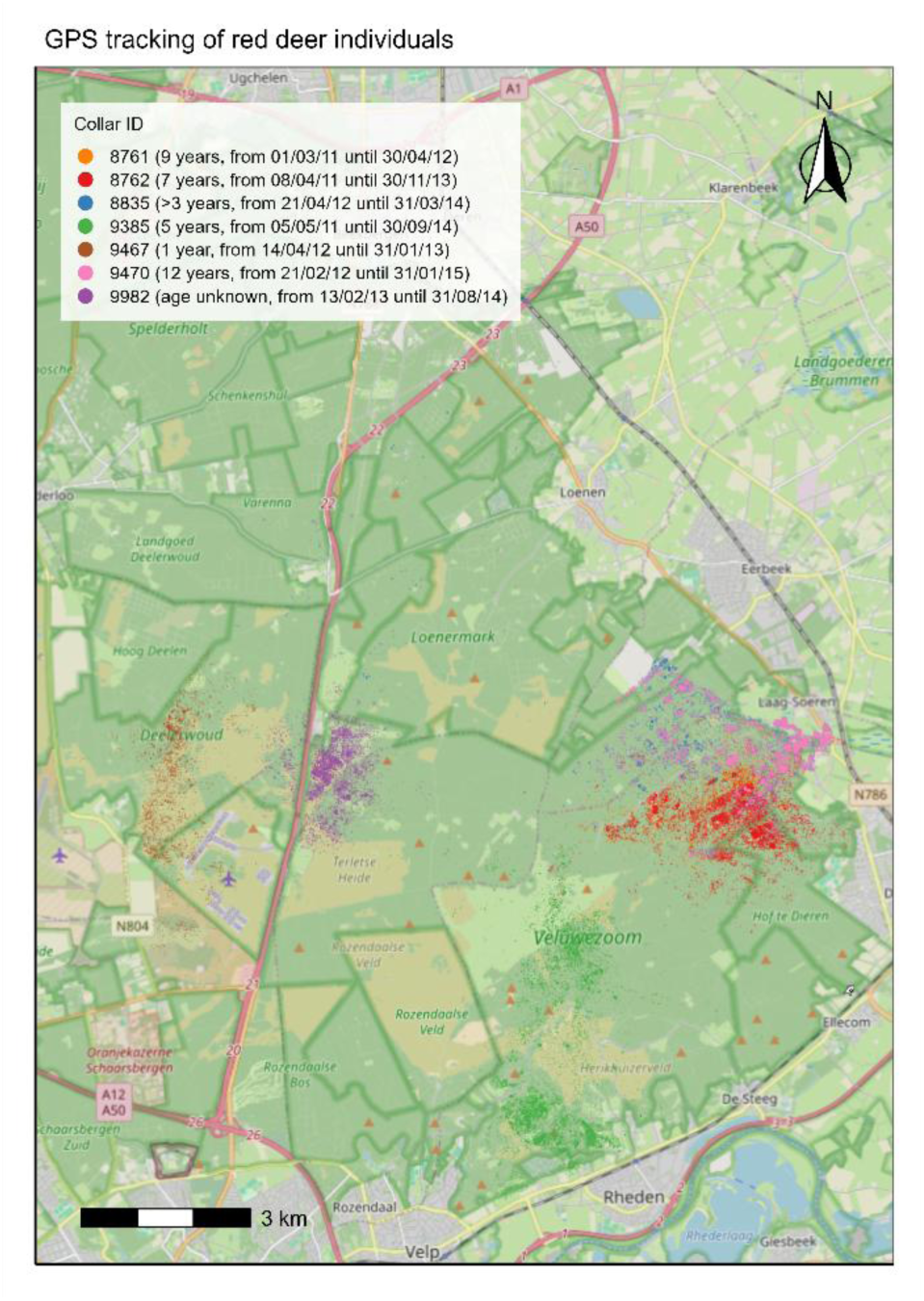
The study area displaying the GPS locations for the studied red deer individuals, for females and males. The legend lists the collar ID for each individual, along with their age and the start and end dates of GPS tracking.

**Table 1.** Sex, age, date of collaring, and running time per individual red deer.

| ID | Sex | Age (years) | Date of blood sampling and GPS collaring | Running time |
| --- | --- | --- | --- | --- |
| 8761 | Male | 9 | 15-02-2011 | 01-05-2011 – 30-04-2012 |
| 8762 | Female | 7 | 24-03-2011 | 01-01-2012 – 31-12-2012 |
| 8835 | Female | 2-3 | 07-04-2012 | 01-01-2013 – 31-12-2013 |
| 9385 | Female | 5 | 21-04-2011 | 01-01-2013 – 31-12-2013 |
| 9467 | Male | 1 | 20-02-2012 | 01-05-2012 – 31-12-2012 |
| 9470 | Male | 12 | 07-02-2012 | 01-01-2013 – 31-12-2013 |
| 9982 | Female | Unknown | 30-01-2013 | 01-03-2013 – 28-02-2014 |

Creating a clean set of locations, obvious outliers of the GPS data - for instance, GPS-fixes within buildings – were removed visually. Then, we calculated monthly home ranges as 100% minimum convex polygons using the “amt” package (Signer et al. 2019) in R version 4.5.1 (R Core Team 2025). Including seasonal variation, 12 consecutive months were selected for each individual. To ensure comparable sampling effort among months, only monthly home ranges based on minimum 500 GPS locations were used. One individual had a total tracking duration of only 9 months, which was still included.

### 2.4 Soil nutrient heterogeneity

For each individual, monthly home ranges were annotated with soil types (Bodemdata.nl n.d.; IUSS Working Group WRB 2022) using ArcGIS pro (version 2.9.5), yielding a total of nineteen soil types across all home ranges. A-priori further analyses, we assigned each soil type a nutrient score based on soil pH, cation exchange capacity (CEC), and soil structure (Table 2). Highly detailed information about the local nutrient contents was not available. Values for pH and CEC were obtained from thematic maps in ArcGIS Online (Esri ArcGIS Online, n.d.; Esri n.d.) and combined with a soil classification map (Bodemdata.nl n.d.). Mean pH and CEC values were calculated per soil type using the Zonal Statistics as Table tool in ArcGIS.

**Table 2.** Soil pH, cation exchange capacity (CEC), and assigned nutrient scores for the 19 soil types in the study area. Mean pH and CEC values were derived from ArcGIS-based thematic maps using zonal statistics. Nutrient scores (1 = relatively nutrient-poor; 3 = relatively nutrient-rich) were assigned based on combined information on pH, CEC, and soil structure. All soils were acidic (pH < 6) with low CEC (< 12), indicating generally nutrient-poor conditions.

| Soil type | pH | CEC | Nutrient score |
| --- | --- | --- | --- |
| Plaggen soils with peat influence; loamy fine sand (Beekeerdgronden) | 5.6 | 10.9 | 3 |
| Weakly developed dune podzol soils; loam-poor, slightly loamy fine sand (Duinvaaggronden) | 4.9 | 7.1 | 1 |
| Peaty topsoil over loamy fine sand (Gooreerdgronden) | 5.4 | 10.2 | 3 |
| Podzol soils on coarse sand (Haarpodzolgronden) | 5.0 | 7.5 | 1 |
| Podzol soils; loam-poor, slightly loamy fine sand (Haarpodzolgronden) | 4.9 | 7.5 | 1 |
| Plaggen soils; anthropogenic, loamy fine sand | 5.5 | 9.6 | 3 |
| Anthropogenic plaggen soils; high organic topsoil; coarse sand (Hoge zwarte enkeerdgronden) | 5.2 | 7.1 | 1 |
| Podzol soils on coarse sand (Holtpodzolgronden) | 4.9 | 7.2 | 1 |
| Podzol soils; loam-poor, slightly loamy fine sand (Holtpodzolgronden) | 5.0 | 7.9 | 2 |
| Podzol soils on loamy fine sand (Holtpodzolgronden) | 4.8 | 7.2 | 1 |
| Podzol soils on coarse sand (Laarpodzolgronden) | 5.4 | 10.3 | 2 |
| Podzol soils; loam-poor and slightly loamy fine sand (Laarpodzolgronden) | 5.4 | 9.4 | 2 |
| Podzol soils on coarse sand (Loopodzolgronden) | 5.0 | 5.2 | 1 |
| Podzol soils; loamy fine sand (Loopodzolgronden) | 5.2 | 8.2 | 2 |
| Peat soils over sand without humus podzol, shallow (<1.2m) (Meerveengronden) | 5.4 | 9.2 | 2 |
| Weak podzolised soils; sandy loam in situ (Ooivaaggronden) | 4.9 | 9.0 | 2 |
| Podzol soils on coarse sand (Veldpodzolgronden) | 5.4 | 9.6 | 2 |
| Podzol soils; slightly loamy fine sand (Veldpodzolgronden) | 5.5 | 11.7 | 3 |
| Weak podzolised soils; loam-poor sandy soils (Vlakvaaggronden) | 5.0 | 7.8 | 1 |

CEC reflects a soil’s capacity to retain and exchange nutrient cations (e.g. Ca, Mg, K), with higher values indicating greater nutrient-holding capacity (e.g. Zajícová & Chuman 2019; Madueke et al. 2021). Soil pH strongly influences CEC by affecting the variable negative charge on soil colloids (Weil & Brady 2016); under more acidic conditions, protonation reduces negative charge and thus cation retention, meaning that a modest increase in pH within the acidic range (e.g. from 4.8 to 5.6) can result in a higher CEC. In addition, soil texture – particularly the proportion of clay, sand, and organic matter – further determine CEC. Clay-rich soils generally exhibit higher CEC due to a greater number of binding sites, whereas in sandy soils CEC is largely dependent on organic matter content (e.g. Zajícová & Chuman 2019; Madueke et al. 2021). Therefore, we used the combination of CEC, pH and soil texture to assign each soil type a nutrient score.

Based on the combined information on pH, CEC, and soil structure, we assigned each soil type in our study area (Table 2) a nutrient score ranging from 1 (relatively nutrient-poor) to 3 (relatively nutrient-rich). Although all soils were acidic (pH < 6) and had low CEC (< 12), indicating overall nutrient-poor conditions in the Veluwe region, they differed in relative nutrient availability, hence the region was suitable for examining our research aim. In addition to variation among soil types, the study area also contained former agricultural fields that represented locally enriched habitat patches and were analysed separately (see prediction iv below).

### 2.5 Statistical analyses

Prior to testing the predictions, we calculated the monthly average nutrient score of both the total home range and the locations used by the red deer within the home range. The average nutrient score of the total home range was calculated as the area-weighted mean nutrient score across all soil types within the home range. The average nutrient score of used locations was calculated as the mean nutrient score weighted by the number of GPS locations recorded on each soil type. For each individual and month, the home range size (ha) was derived from the 100% minimum convex polygon estimates based on GPS locations, and linked to the corresponding average nutrient scores of both the total home range and the used locations.

Then, first, to test whether individuals in more nutrient-poor areas maintained larger home ranges (prediction i), we examined the relationship between nutrient score and monthly home range size for the total home range size and used locations within the home range. We used linear mixed-effects models (Bates et al. 2015) with log-transformed home range size as the response variable and average nutrient score as a fixed effect. Sex and month were included as additional fixed effects to account for sex-specific and seasonal variation, and individual identity (CollarID) was included as a random intercept to account for repeated monthly observations of the same individuals. Separate models were fitted for nutrient scores of the total home range and used locations.

Second, to test whether individuals used the most nutrient-rich areas within their home ranges (prediction ii), we used a linear mixed-effects model (Bates et al. 2015) with average nutrient score as the response variable, and sex, home-range type (used locations versus total home range), and their interaction as fixed effects. Month was included as an additional fixed effect to account for seasonal variation, and individual identity (CollarID) was included as a random intercept to account for repeated monthly observations of the same individuals. Significance of fixed effects was assessed using analysis of variance, and when relevant, post hoc pairwise comparisons were performed using estimated marginal means with the emmeans package (Lenth et al. 2019).

Third, to test whether individuals foraged more on soil types with higher nutrient scores (prediction iii), we compared the observed use of soil types with their expected use based on availability within their home range. For each individual and month combination, the number of GPS locations recorded on each soil type was used as a measure of observed use. The expected use was calculated as the proportion of each soil type within the home range multiplied by the total number of locations. Selection for each soil type was then quantified as the ratio of observed to expected use, where values greater than 1 indicate selection and values less than 1 indicate avoidance. We performed G-tests of goodness-of-fit for each soil type within each individual-month combination. For each soil type, we used a one-versus-rest approach, in which the observed number of locations on a given soil type was compared to the combined observations on all the other soil types. To account for multiple testing across soil types within each individual-month combination, p-values were adjusted using the Benjamini-Hochberg false discovery rate procedure (Benjamini & Hochberg 1995). Analyses were restricted to cases with at least 500 observations and non-zero expected values.

Last, to test whether individuals selected locally enriched patches associated with former agricultural land use more than surrounding areas (prediction iv), we compared the observed use of these patches with their expected use based on availability within the monthly home range. Former agricultural patches, varying in size between 0.3 and 65 hectares, were defined using a spatial polygon layer based on the knowledge of local rangers and historical aerial photographs, and GPS locations were classified as either inside or outside these patches. Then, we used G-tests of goodness-of-fit for each individual-month combination, with p-values adjusted using the Benjamini-Hochberg false discovery rate procedure to determine significance for each month (Benjamini & Hochberg 1995).

## 3. Results

### 3.1 Home range size versus nutrient richness

The monthly home ranges varied between 172 and 5,131 hectares, with an average of 827 ± 867 hectares (mean ± SD). We found no evidence that individuals in nutrient-poor areas maintained larger home ranges (Fig. 2). After accounting for repeated observations of individuals, sex, and month, home range size was not significantly related to the average nutrient score of either the total home range (Fig. 2; LMM, df = 36.9, F = 1.99, p = 0.167) or the used locations (Fig. 2; LMM, df = 50.2, F = 2.37, p = 0.130).

**Figure 2.**
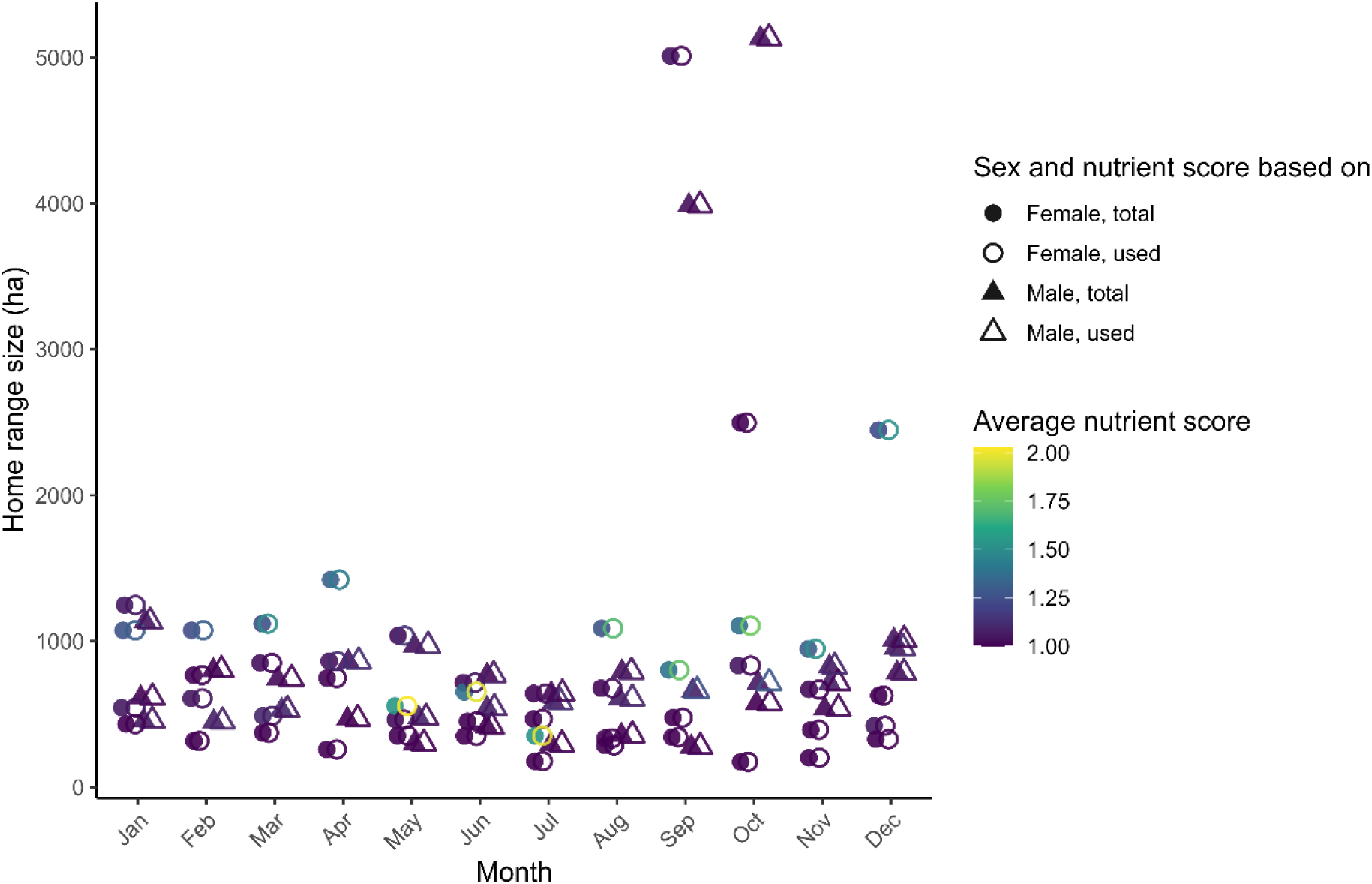
The monthly home ranges sizes of individual red deer, distinguished by used and total home range. The colour indicate average nutrient score, with darker colours reflecting lower nutrient scores.

### 3.2 Use of nutrient-rich home-range patches

Overall, average nutrient scores were higher for used locations than for the total home range, indicating that red deer predominantly used relatively nutrient-rich areas within their home ranges (Fig. 3). This pattern was more pronounced in females than in males.

**Figure 3.**
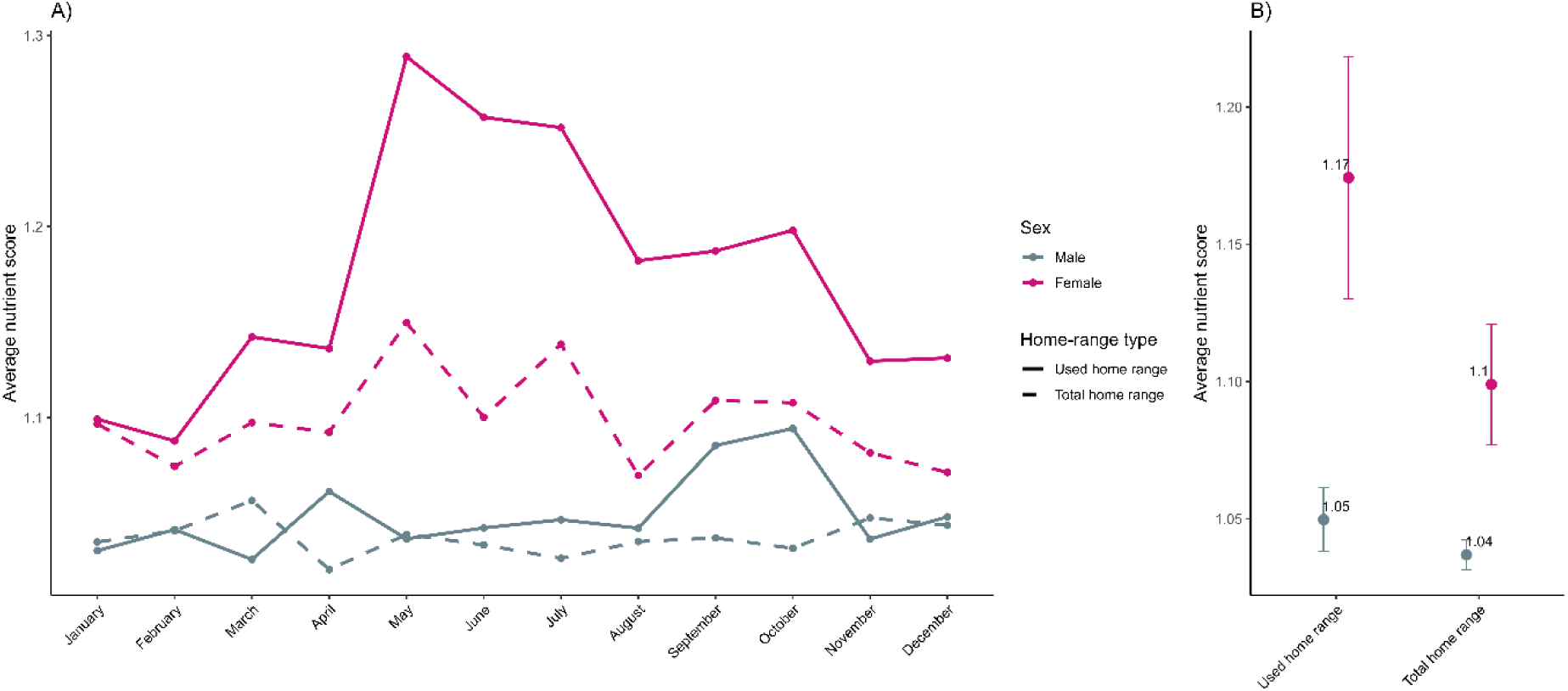
Monthly variation in average nutrient scores of reed deer home ranges (A), distinguished by used (solid lines) and total (dashed lines) home ranges, and separated by sex (colours), and the overall mean nutrient scores (± SE) for each category (B).

The linear mixed-effects model showed a significant effect of home-range type (i.e. total home range versus used locations) on nutrient score (LMM, df = 1,145, F = 11.57, p < 0.001), as well as a significant interaction between sex and home-range type (LMM, df = 1,145, F = 4.39, p = 0.038). There was no significant main effect of sex (LMM, df = 1,145, F = 0.54, p = 0.464), and month did not significantly affect the nutrient score of the home-range (LMM, df = 11,145, F = 0.91, p = 0.533). Post hoc comparisons showed that females used areas with higher nutrient scores than expected based on availability (estimate = 0.075, p < 0.001), whereas no significant difference between used and total home ranges was found for males (estimate = 0.013, p = 0.58).

### 3.3 Selection for nutrient-rich soil types

We found that individuals used some soil types either more (ratio > 1) or less (ratio < 1) frequently than expected based on the availability of each soil type within the home range (Fig. 4; Table S1.1). For example, proto-Carbic podzol soils (nutrient score 2) were used substantially more often than expected (median observed/expected ratio = 5.93, IQR = 4.32-7.08), and anthropogenic plaggen soils with loamy fine sand (nutrient score 3) were also more frequently selected (median = 2.48, IQR = 2.20–3.18). In contrast, weak podzolised soils (nutrient score 1) and peaty topsoil over loamy fine sand (nutrient score 3) were used less frequently than expected, with median ratios close to zero, indicating strong avoidance despite their availability within the home range.

**Figure 4.**
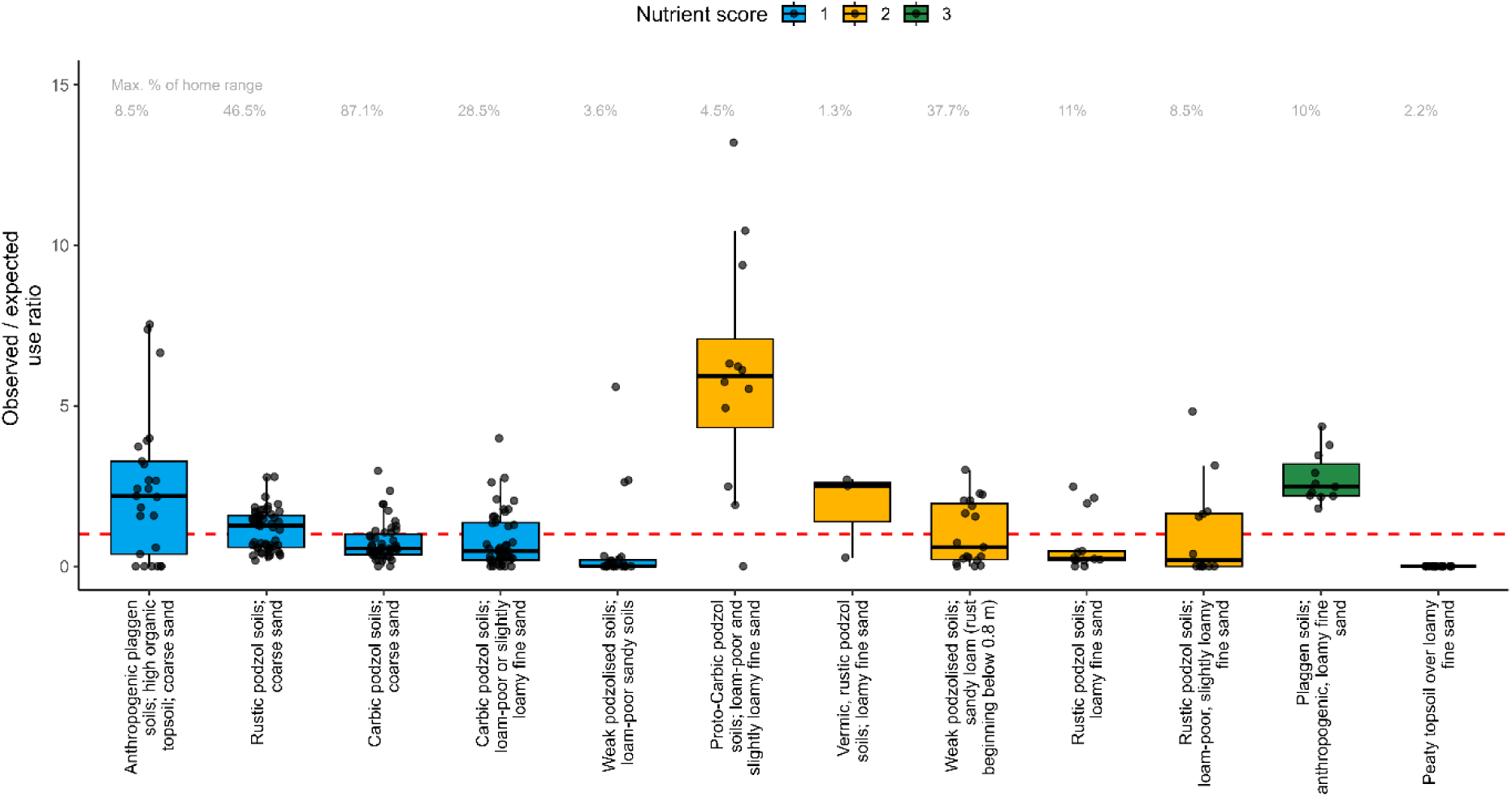
The significant (adjusted p-value < 0.05) ratio values per soil type. Values > 1 indicate selection and values < 1 indicate avoidance.

Selection was not consistently aligned with nutrient score (Fig. 4). Although some high-scoring soils were selected, others were avoided. Similarly, intermediate- and low-scoring soils showed contrasting patterns: proto-Carbic podzol soils (score 2) were strongly selected, whereas rustic podzol soils with loamy fine sand (score 2) were avoided (median = 0.22), and weak podzolised soils with loam-poor sandy soils (score 1) were strongly avoided (median = 0.03), whereas anthropogenic plaggen soils with high organic topsoil and coarse sand (score 1) were selected (median = 2.19).

### 3.4 Selection for former agricultural patches

Former agricultural patches were, on average, used more frequently than expected based on their availability within monthly home ranges (median observed/expected ratio = 1.31, IQR = 0.22–4.29; Fig. 5). G-tests of goodness-of-fit further showed that patch use was often non-random relative to availability, with 69 of 139 individual–month combinations remaining significant after Benjamini-Hochberg correction for multiple testing (*p* < 0.05; Table S1.2). Selection was the strongest in winter and spring, and least in late summer and autumn (Fig. 5).

**Figure 5.**
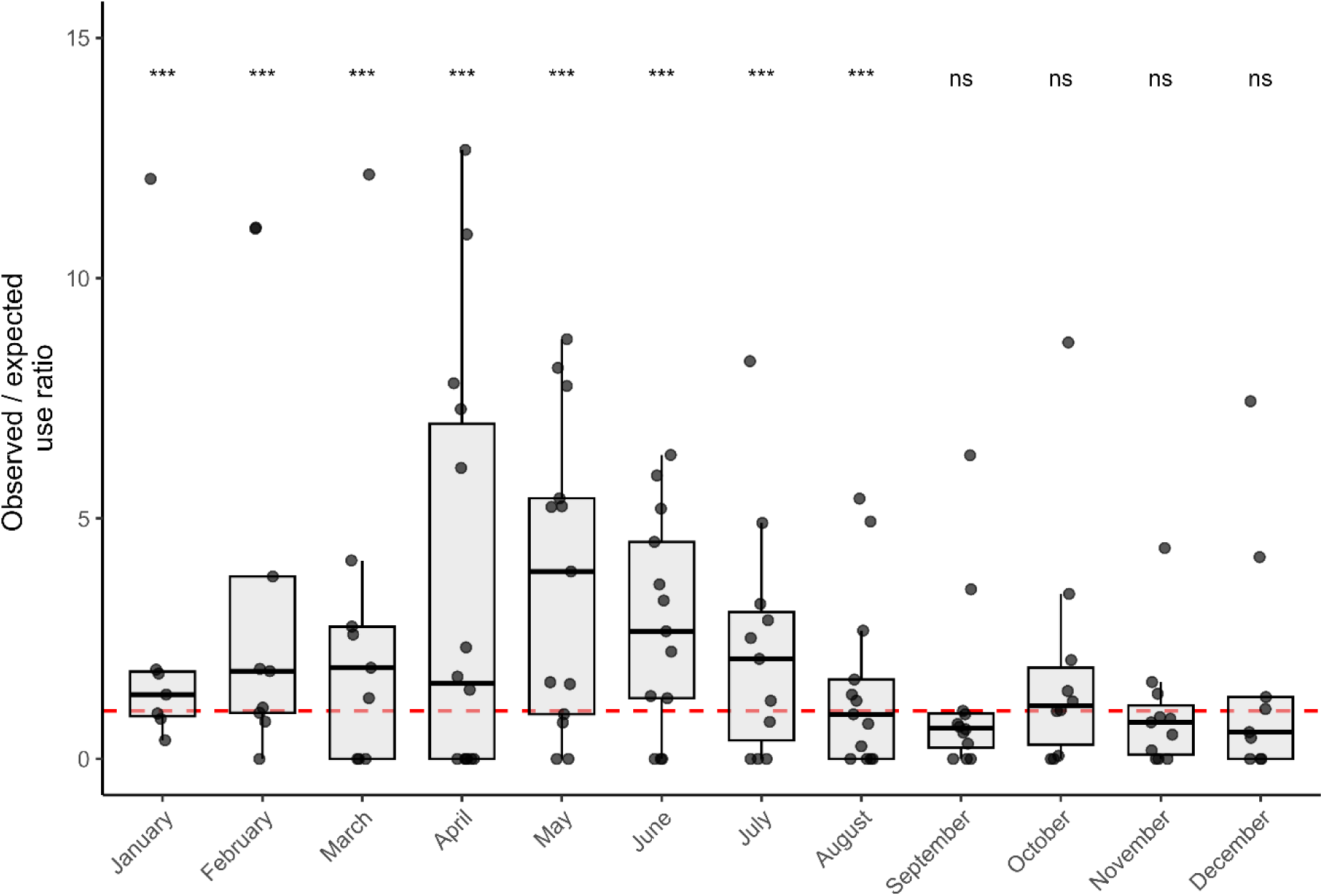
The significant (adjusted p-value < 0.05) ratio values for former agricultural patches per month. Values > 1 indicate selection and values < 1 indicate avoidance.

## 4. Discussion

We investigated how nutrient heterogeneity, as reflected by soil types and former agricultural patches, shaped habitat use and selection by a large herbivore. Our results partially support the hypothesis that spatial variation in soil-derived nutrient availability structures habitat selection by large herbivores under generally nutrient-poor conditions. Red deer used the relatively nutrient-rich patches and locally enriched former agricultural patches within their home ranges more extensively than expected based on availability in their home range. However, selection did not follow a consistent gradient in nutrient score among soil types. Instead, selection was strongest for specific soil types: anthropogenic plaggen soils with loamy fine sand, which had a high nutrient score, and proto-Carbic podzol soils with loam-poor and slightly loamy fine sand, which may reflect favourable physical habitat conditions such as relatively dry and accessible terrain (IUSS Working Group WRB 2022). Together, these findings suggest that nutrient availability influences habitat selection primarily through fine-scale and context-dependent processes, rather than through broad-scale gradients in soil-derived nutrient availability.

At the scale of the home range, individuals showed a tendency to use relatively nutrient-rich areas within the habitat available to them. This pattern indicates that red deer can exploit spatial variation in resource quality within their accessible habitat, consistent with selective foraging theory to optimise nutrient intake (Raubenheimer et al. 2009; Raubenheimer & Simpson 2016). The stronger effect observed in females suggests that nutritional requirements may differ between sexes (e.g. Johan 2005; Maklakov et al. 2008), potentially reflecting higher energetic demand during the reproduction season (Flueck et al. 2012; Ceacero et al. 2015; Wenting et al. 2023). Such sex-specific responses are consistent with studies showing that females often exhibit more selective habitat use when resource quality is limiting (e.g. Clutton-Brock et al. 1987; Alves et al. 2014).

Individuals in more nutrient-poor areas did not maintain larger home ranges. After accounting for repeated observations of individuals, sex, and seasonal variation, home range size was not significantly related to the nutrient score of either the total home range or the used locations within it. This suggests that monthly home range size is not primarily determined by broad-scale variation in soil-derived nutrient availability. Instead, home range size is likely shaped by multiple other factors, including social interactions (e.g. Krojerová-Prokešová et al. 2010; Albery et al. 2022; Jarnemo et al. 2023), reproductive activities (e.g. Parker et al. 2009; Pagon et al. 2017), landscape configuration (e.g. Bevanda et al. 2015; Jarnemo et al. 2023), and habitat diversity within home ranges (Ullmann et al. 2018). These findings highlight that home range size cannot be interpreted as a straightforward proxy for resource scarcity.

Former agricultural patches were used more frequently than expected based on their availability, indicating that red deer do respond to localised nutrient hotspots. Selection was strongest in late winter and early spring, suggesting that nutrient-rich patches may be particularly important during periods of increased nutritional demand or reduced resource availability (e.g. Grace & Wilson 2002; Grace et al. 2008). However, patch selection varied among individuals and months, with both strong selection and avoidance observed. This variability suggests that the use of enriched patches is influenced by additional factors such as seasonal changes in resource availability, individual behaviour, local landscape context, and competition with other species (e.g. Pérez-Barbería et al. 2013; Fattebert et al. 2019; Jarnemo et al. 2023). Also, it could suggest that the used nutrient scores and former agricultural patches do not precisely reflect nutrient availability in the soil nor in the vegetation, that we consider a potential limitation of our study. We were not able to correct for this due to unavailability of more precise nutrient data in the landscape. Seasonal changes in vegetation quality and phenology are likely to influence the relative importance of different habitat components (e.g. Wiegand et al. 2008; Liang & Schwartz 2009), while differences among individuals may reflect variation in behaviour, condition, or life-history stage (e.g. Jarnemo et al. 2023).

The importance of historically enriched patches, such as the former agricultural patches in our study, highlights the lasting ecological effects of past land use, even in landscapes that are managed to promote natural processes. Our findings therefore have implications for understanding the role of large herbivores in nutrient dynamics and ecosystem functioning. By selectively using nutrient-rich patches, herbivores may contribute to the redistribution of nutrients across the landscape (e.g. Kie et al. 2005; Ganskopp & Bohnert 2009; Riesch et al. 2022), potentially reinforcing or modifying existing spatial heterogeneity. In the absence of large predators, such as in the Veluwe at the time that our GPS data was collected, herbivore movements may be less constrained by predation risk (e.g. Theuerkauf & Rouys 2008; Vanderlocht et al. 2025; Van Beeck Calkoen et al. 2026), potentially amplifying their influence on spatial patterns of resource use. As a result, selective use of nutrient-rich patches may not only reflect underlying heterogeneity, but also contribute to its maintenance or reinforcement through processes such as dung deposition and differential grazing pressure (e.g. Catorci et al. 2016; Hjermann et al. 2025). In systems affected by long-term N deposition and nutrient imbalances, such processes may considerably influence vegetation composition, competitive interactions, and broader ecosystem functioning.

These findings may also be relevant for restoration and management interventions aimed at improving nutrient availability in nutrient-depleted ecosystems. For example, silicate rock powder has been recently – not during the timeframe the red deer were GPS-tracked – applied in parts of the Veluwe to counteract soil acidification and replenish depleted base cations (e.g. Van Der Bauwhede et al. 2025; Harmsen et al. 2026). Such measures could, in principle, increase habitat quality and attractiveness for herbivores by improving forage conditions. However, in the same region as our study area, no evidence has been found that foraging ungulates – including red deer – were attracted to patches treated with silicate rock powder within three years after application (Heurman et al. 2026). Together, these and our findings suggest that herbivore responses may depend not only on nutrient enrichment itself, but also on whether enrichment is sufficiently strong, persistent, and ecologically expressed to alter forage quality at scales relevant for foraging decisions.

A key challenge in linking nutrient availability to animal movement is that soil-derived indicators are only indirect proxies of the expected resource quality (Moore et al. 2020), rather than direct measures of forage quality or nutrient intake. This may partly explain why selection did not consistently relate to nutrient scores assigned to the soil types but is likely to be also mediated by vegetation composition, phenology, and local environmental conditions (e.g. Yamauchi & Yamamura 2004; Van der Waal et al. 2011). Our use of monthly minimum convex polygons defined availability at the home-range scale, allowing us to contrast broad accessible habitat with the locations actually used. The consistent selection of fine-scale nutrient-rich or easily accessible patches (in terms of habitat types) indicates that localised heterogeneity in resource quality and habitat structure plays a meaningful role in shaping habitat use by large herbivores.

## 5. Conclusion

In conclusion, our study provides evidence of selective feeding for nutrients in large herbivores that is partially shape their habitat selection. This effect was not expressed as a simple response to broad-scale soil heterogeneity, but emerged through the disproportional use of fine-scale nutrient-rich patches. Such behaviour may have far-reaching consequences for nutrient redistribution across the landscape and other zoogeochemical processes, with the potential to reinforce, maintain, or alter spatial heterogeneity in resource availability and thereby influence wider ecosystem processes.

## Author contributions

EW, MB and CV led the conceptualisation of this study, with the help of ESB, LS and AK. HL, DRL and RV collected the GPS-data. EW, MB, CV and AK performed the analyses, with assistance of LS and ESB. EW led the writing of the manuscript. All authors contributed critically to the drafts and gave final approval for publication.

## Acknowledgements

We thank everyone involved in the GPS tracking of the red deer, particularly Paul Jansen, Edgar Strikkeling and Frank Theunissen. EW is funded by the research programme Rubicon which is financed by the Dutch Research Council (NWO) under the grant https://doi.org/10.61686/WVSQA32249.

## Data Accessibilty Statement

All the data and supplementary materials used for this manuscript is accessible via Figshare: https:doi.org/10.6084/m9.figshare.33270870.

## Conflict of interest statement

No actual or potential conflicts of interest are declared by the authors.

## Notes

https:doi.org/10.6084/m9.figshare.33270870

### Competing Interest Statement

The authors have declared no competing interest.

